# Contrasting the open access dissemination of COVID-19 and SDG research

**DOI:** 10.1101/2023.05.18.541286

**Authors:** Vincent Larivière, Isabel Basson, Jocalyn P. Clark

## Abstract

This paper examines the extent to which research has been published open access in response to two global threats: COVID-19 and the Sustainable Development Goals (SDGs), including climate change. We compare the accessibility of COVID-19 content versus SDG literature using the Dimensions database between 2000 and 2021, classifying each publication as gold open access, green, bronze, hybrid, or closed. We found that 79.9% of COVID-19 research papers published between January 2020 and December 2021 was open access, with 39.0% published with gold open access licenses. In contrast, just 55.7% of SDG papers were open access in the same time period, with only 36.0% published with gold open access licenses. Papers related to the climate emergency overall had the second-lowest level of open access at just 55.5%. Papers published by the largest for-profit publishers that committed to both the SDG Publishers Compact and climate actions were not predominantly published open access. The paper highlights the need for continued efforts to promote open access publishing to facilitate scientific research and technological development to address global challenges.

**One-Sentence Summary:** In contrast to COVID-19 papers, research on UN Sustainable Development Goals including the climate emergency have not been made open access by leading global science publishers despite their corporate commitments to sustainability and climate action.

## Collective efforts against global threats

Scientific research and technological development are key to solving global challenges. Such importance was clearly highlighted by the rapid production of knowledge for the COVID-19 pandemic response, which required both concerted basic scientific efforts across all fields and the development of new health technologies including vaccines, along with sizeable governmental investments. Within a month of the first reported coronavirus case, more than 160 funders, research institutions, and publishers committed to make such knowledge widely available. The pledge, initiated by the Wellcome Trust in January 2020, committed signatories to share rapidly and openly the research papers and data relevant to the outbreak, in order to inform the global health response and save lives. [1] Open access publishing increases visibility, discoverability, and impact by providing immediate and unlimited access to scientific research. Such a collective commitment to COVID-19 data sharing and open access is remarkable, and begs the question of how such researchers and publishers have responded to other global threats. For example, in 2015, the 154 member states of the United Nations (UN) selected 17 Sustainable Development Goals (SDGs) as part of the 2030 agenda to guide and stimulate global action in areas of critical importance for humanity and the planet. Among these is a commitment to accelerated action on the climate emergency, considered by many as the greatest threat the world has ever faced [3] and in its final crucial decade to change course on devastating health and environmental risks. [4] In recognition, science publishers, including the Big 5 publishers that together represent over 50% market share of a global $28 billion annual publishing industry, [5] agreed to the SDG Publishers Compact. This commits the 270 signatories—which include publishers Elsevier, Sage, Springer Nature, Taylor & Francis, and Wiley—to develop sustainable practices and actively promote content on sustainability, justice, and safeguarding and strengthening the environment. [6]

The world’s largest science publisher, Elsevier, furthermore has specific commitments to climate change explicitly acknowledging their role in supporting the “scholarly dialogue on climate change and its impacts though unique content, data and products” and ensuring that “research is communicated effectively, grounded in real world examples, and acted upon by the right people, [as] one of the greatest challenges we’re facing.”[7] Similarly, the other two largest publishers, Springer-Nature [8] and Wiley [9], have recognised their critical role in promoting climate action and research.

In this context of commitments, we examine the extent to which research has been published open access, comparing accessibility of Coronavirus/COVID-19 content vs SDG, including climate change, literature. Using the Dimensions database of scientific research (www.dimensions.ai) from January 2000 to December 2021 we identified and compared Coronavirus and COVID research publications (including a subset of COVID-19 papers) with research publications classified as SDG related (including a subset of climate related papers), classifying each as to its open access status, defined here in declining levels of openness: gold (immediate and permanent access, author retains copyright and permissions), green (authors self-archive, publisher retains rights), bronze (article is made available but cannot be redistributed or reused), hybrid (published open access in a subscription journal), or not (see supplemental material for full Materials and Methods).

## Not so open science

404,541 COVID-19 research papers were published between Jan 2020 and December 2021, of which 79.9% were open access. Overall, papers on any coronavirus amounted to 417,822 between 2000 and 2021, of which 79.8% are open access including 38.4% gold open access (*Figure 1*). Just 39.0% of the COVID-19 papers were published with gold open access licenses allowing full copyright and re-use allowances, as was required by the Wellcome data sharing pledge [1]. 26.2% of COVID-19 research publications were made available via so-called bronze OA, where a paywall is removed by the publisher but allows them to retain copyright and commercial rights and no obligation to maintain access for any period of time for non-paying readers. Nevertheless, 3 years after COVID-19 was declared a public health emergency of international concern by the World Health Organization, publishers have continued to make COVID-19 content freely available via various open access routes. This policy is at least partially responsible for the unprecedented volume science publishers have received of article submissions, publications, and market growth, which is reported to be 8.1% for 2020, three times larger than the average annual absolute growth from 2013 to 2019. [5,10]

**Figure 1.**
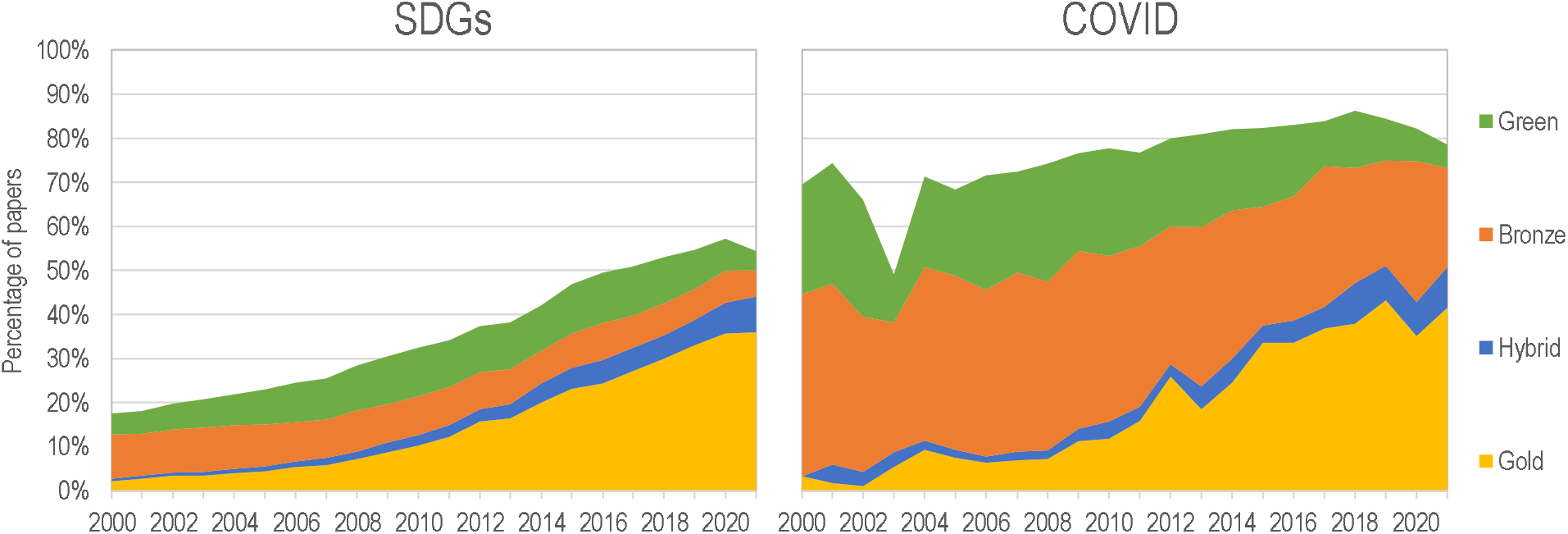
Percentage of open access for SDG-related and COVID-related research papers, by access type, 2000-2021.

In contrast, 803,152 distinct papers on an SDG were published between Jan 2020 and the end of 2021, and just 55.7% of these were open access as of June 2022. 36% of the SDG papers were gold open access. Bronze, green, and hybrid open access licenses each accounted for less than 10% of papers. Levels of open access of relevant research varied by SDG, but none attained levels of COVID-19 (*Figure 2*). Most SDGs have openness rates between 55% and 67%, except for affordable and clean energy (SDG 7) for which a minority of papers are freely available. Among these, papers related to climate change ranks second lowest in in terms of open access, at just 55.5% overall. A third of research papers (33.8%) on the climate emergency are published with a gold open access license allowing reuse and author retention of copyright.

**Figure 2.**
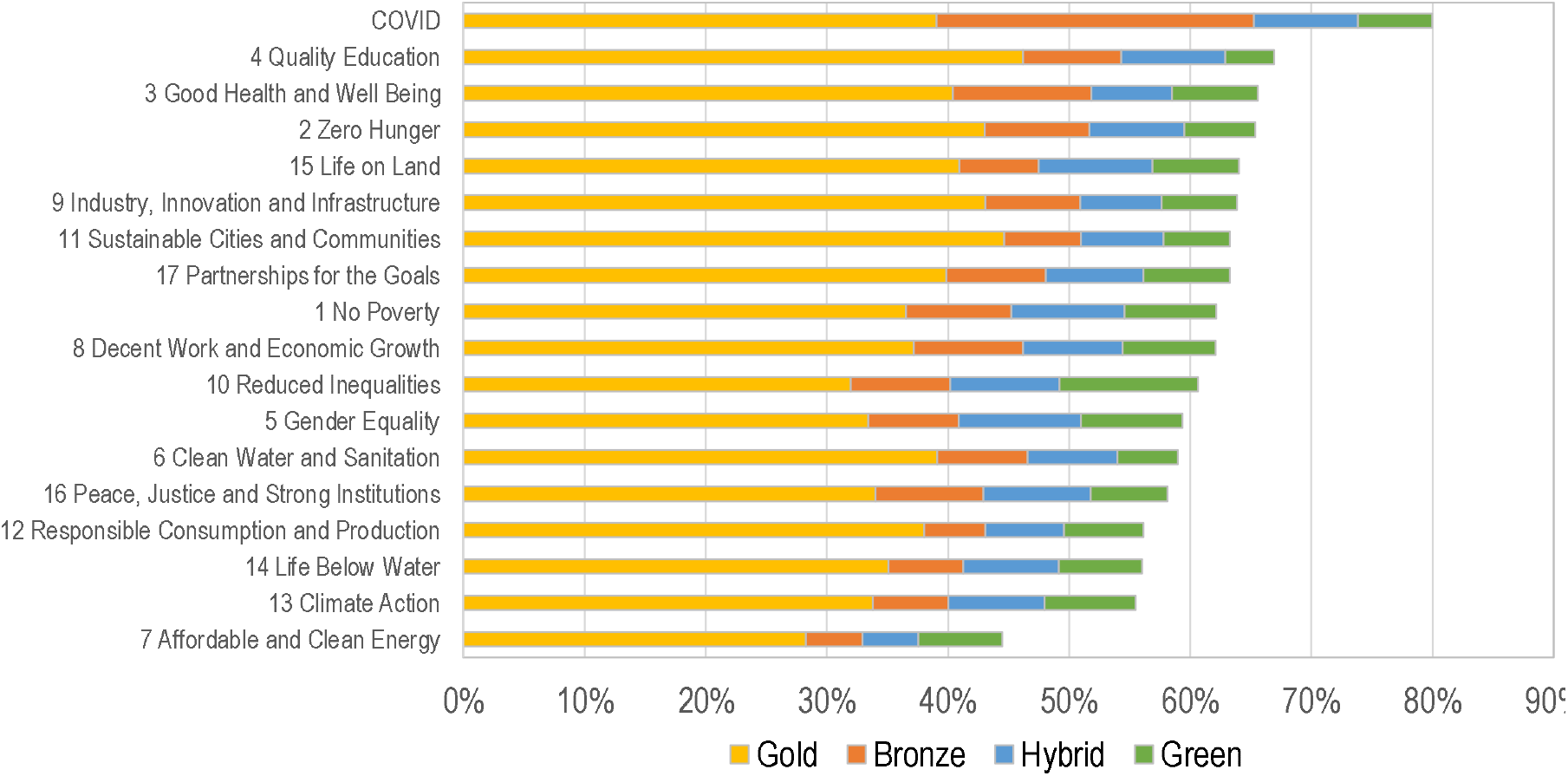
Number of papers by SDG and for COVID-related research, by access type, 2017-2021. Dimensions.ai database.

**Figure 3.**
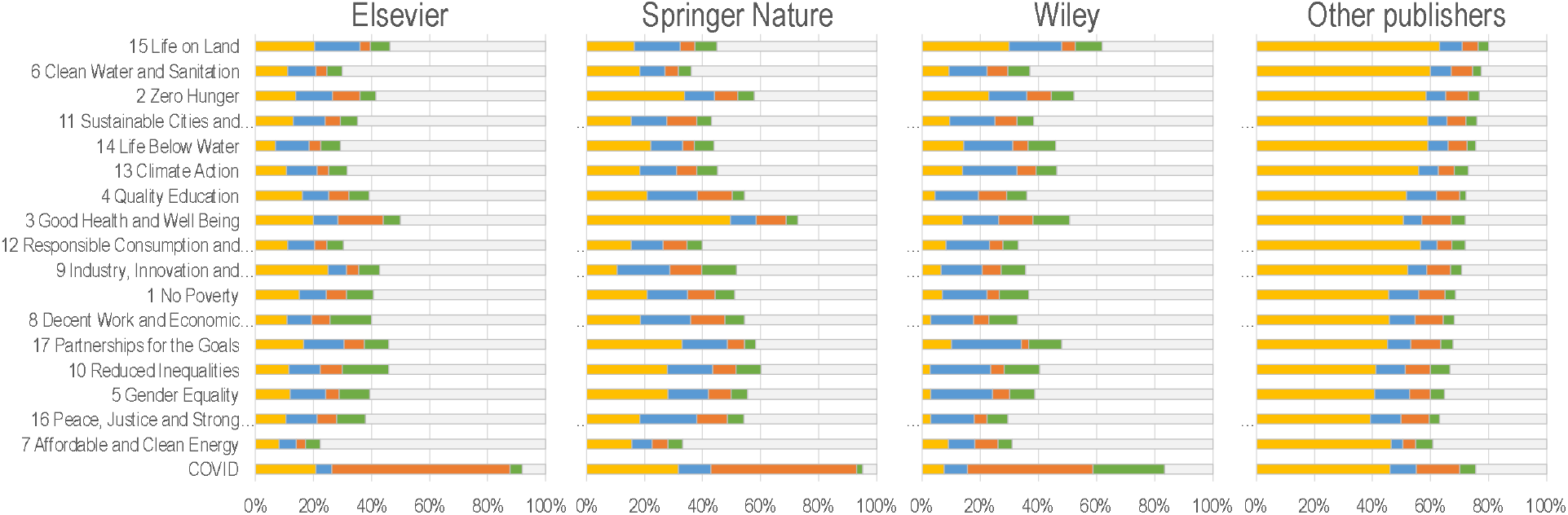
Percentage of open access (gold, hybrid, bronze and green) papers, for sustainable development goal research and COVID-related research, by publisher, 2015-2021.

For work published in the largest publishers that committed to both the SDG Publishers Compact and climate actions, related papers are not predominantly published open access. For example, among all SDG research papers published by Elsevier, no SDG has more than 50% of its research papers disseminated via open access publishing between 2017 and 2021. These levels range from 22.2% for papers on affordable and clean energy (SDG7) to 49.9% of those related to health and well-being (SDG3). Percentages are slightly higher for papers published by Springer-Nature or Wiley, with respectively ten and three SDGs having more than 50% of the related papers open access. For the climate emergency (SDG13), none of the three top publishers made more than 50% of the papers on this topic that they published open access. In contrast, the top 3 largest publishers have made almost all COVID-19 papers by June 2022 openly available to the research community and the general public: 92.1% of those published by Elsevier, 95.0% by Springer Nature, and 83.4% by Wiley.

## Publishing paradoxes

What does this say about science publishing in the midst of global threats? COVID-19 seemed to spur collective action on the part of the research and publishing communities, which had no precedent but nonetheless demonstrated the ability to rapidly and openly disseminate crucial knowledge. Publishing executives and editors may have felt a moral obligation to contribute to collaborative efforts when global solidarity was required to overcome a novel infectious disease threat. An assessment of the Wellcome pledge similarly found that about half of COVID-19 papers published among its signatories were gold open access. [11] In contrast, despite the enormity and importance of the challenge at stake with the climate emergency – the very survival of the planet and the human race – the vast majority of papers published on the topic are not open access. This inconsistency was particularly acute for the top 3 largest publishers, which are home to the world’s most prestigious and highly visible journals. The contrast between openness of research related to the COVID-19 pandemic but not to the priority global development goals or the climate emergency raises a critical tension in the scholarly communication field. It is a worrying contradiction in how and what knowledge is published and disseminated that health advocates, funders, and institutional leaders must demand more attention to.

A main element to consider is the role of bronze OA, which is a way to make content freely available without giving up commercial rights. It has the appearance of being and fulfilling “open access” commitments, whilst allowing publishers to retain copyright and block redistribution or reuse. We observe that for COVID-19 research, the bronze OA route was utilised by the largest 3 publishers. But for the other publishers, gold open access was predominantly provided. Some of this may be due to the choice of authors rather than publishers, but is unlikely to be a sizeable share of papers given the promise of many publishers to ensure COVID-19 content is made available. Among the top 3 largest publishers, therefore, access to COVID papers assigned bronze OA can be withdrawn at any time. Retention of copyright and reuse may also serve a commercial strategy to preserve dominant market position and historical huge profit margins. [5, 12] The extent to which publishers can market and sell gold open access publishing, which is often facilitated via article publication charges paid by the author, institution or funder, is evident in the SDG literature. In the Dimensions database, about 18.2% overall of research is published this way, compared with 12.3% for bronze OA [13]. A large proportion of COVID-19 publications, produced at volumes never seen before in the scientific literature, are thus owned by publishers rather than by their authors. The research asset created by the extraordinary collective response of scientists and funders to COVID-19 worldwide is not a wholly public one.

## The sustainability challenge

The importance of scientific knowledge and the global impacts of health threats are at their highest levels of public consciousness. Therefore, the general public and readers alike will increasingly expect scientific institutions to uphold higher-than-average moral standards and to be driven by common good principles rather than personal advancement. The SDG agenda is multi-faceted but no less important than COVID-19 as a global health challenge, and like climate change is not a mere scientific concern, but a social one; requiring concerted, collective global efforts to tackle and implement the blueprint for current and future peace and prosperity for people and the planet that was envisioned when member states agreed to the sustainable development priorities. Equally, publishers who agreed to sustainable development and climate action must be held to account for their commitments and plans. Facilitating SDG literature, including research relevant to the climate emergency, to be open to all would inform researchers, policymakers, and the public. Knowledge is critical to understand what is at stake, advance scientific outcomes, and help progress towards sustainability and climate goals, most of which have been severely derailed by the COVID-19 pandemic. [14]

## Author contributions

Conceptualization: VL, JC

Data Curation: VL

Funding acquisition: VL

Investigation: VL, IB

Methodology: VL

Project administration: VL

Resources: VL

Supervision: VL

Visualization: VL

Writing - Original Draft: VL, IB, JC

Writing - Review & Editing: VL, IB, JC

## Supplementary Materials

Figs. S1 to S2

## Supplementary Materials

Using the Dimensions database from January 2000 to June 2022, we created two datasets: one of COVID-related research (including research on coronaviruses in general) and one related to SDGs. The COVID dataset was created using the COVID*, SARS-COV-2 and Coronavirus* search strings, which has been the standard query to retrieve papers on the topic.^1^ This query retrieved 417,822 research papers between 2000 and 2021; most of which have been published since 2020 (96.8%). The definition of research related to SDGs remains more challenging, as some of the SDGs remain poorly defined.^2^ However, recent advances in natural language processing and machine learning led to the creation of a validated dataset of SDG related research, which we have leveraged for this analysis.^3^ The SDG-related dataset consists of 4,636,992 paper/SDG combinations (as papers can be associated with more than one SDG). The smallest SDG from a number of paper point of view is SDG 17 (Partnerships for the Goals) with 9,150 papers, followed SDG 9 (Industry, Innovation and Infrastructure), which contains 14,509. At the other end of the spectrum, SDG 3 (Good Health and Well Being) has the highest numbers of papers (1,178,622), followed by SDG 7 (Affordable and Clean Energy), with 1,047,233 papers (Fig. S1). Open access of papers was determined by Unpaywall, as embedded in the Dimensions.ai database. Following Piwowar et al.,^4^ open access status was divided into four mutually exclusive category: gold open access (paper published in a fully open access journal— with or without APCs), hybrid open access (paper published with an open license in a subscription journal), bronze open access (paper freely available on the journal website, but without a clear license), and green open access (papers solely available in a repository). It is worth recalling that the open access status is determined as of now (or a couple months ago) and does not represent the availability at the time of publication.

**Supplementary Figure S1.**
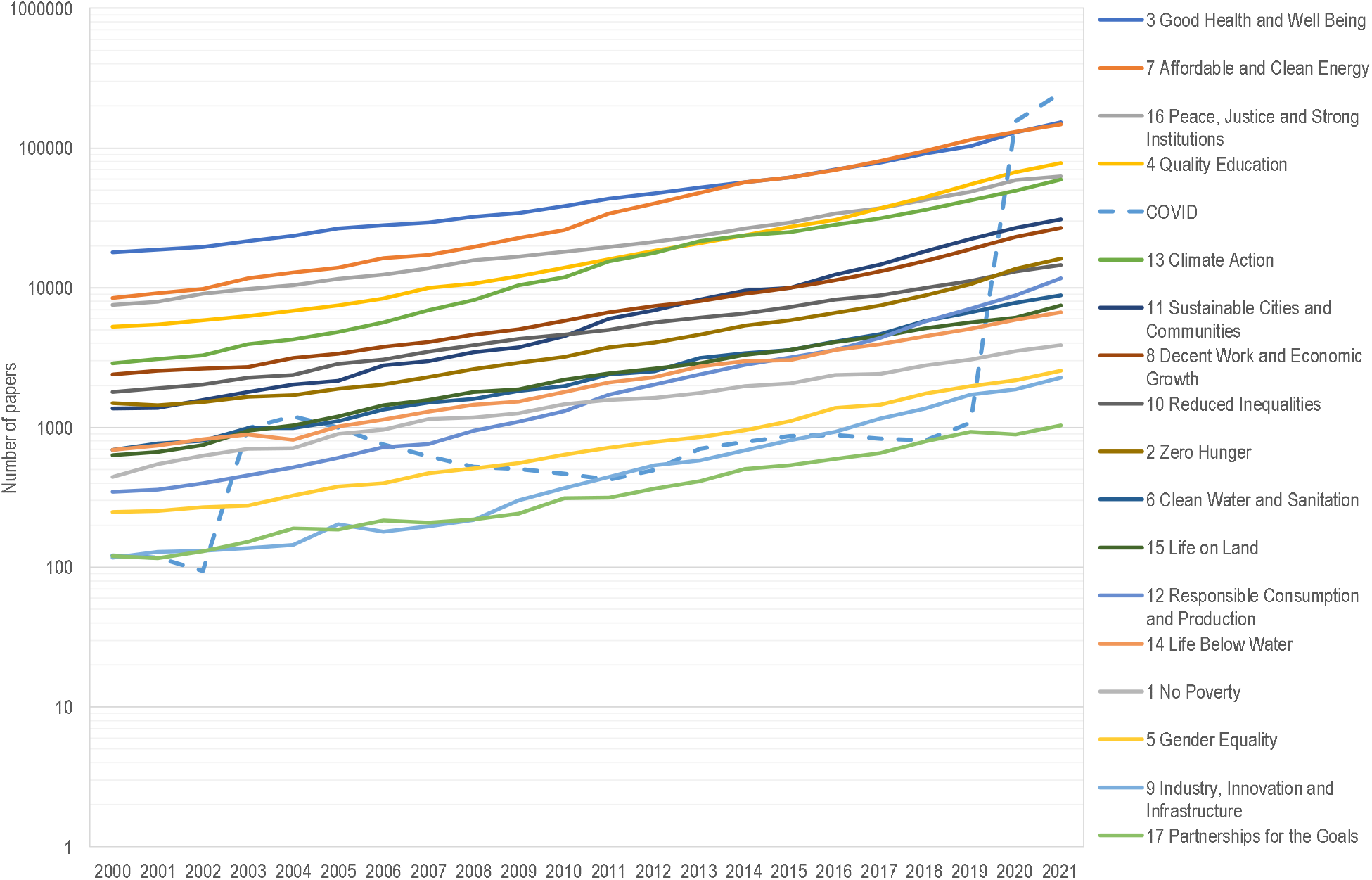
Number of papers by SDG and for COVID-related research, 2000-2021. Dimensions.ai database.

**Supplementary Figure S2.**
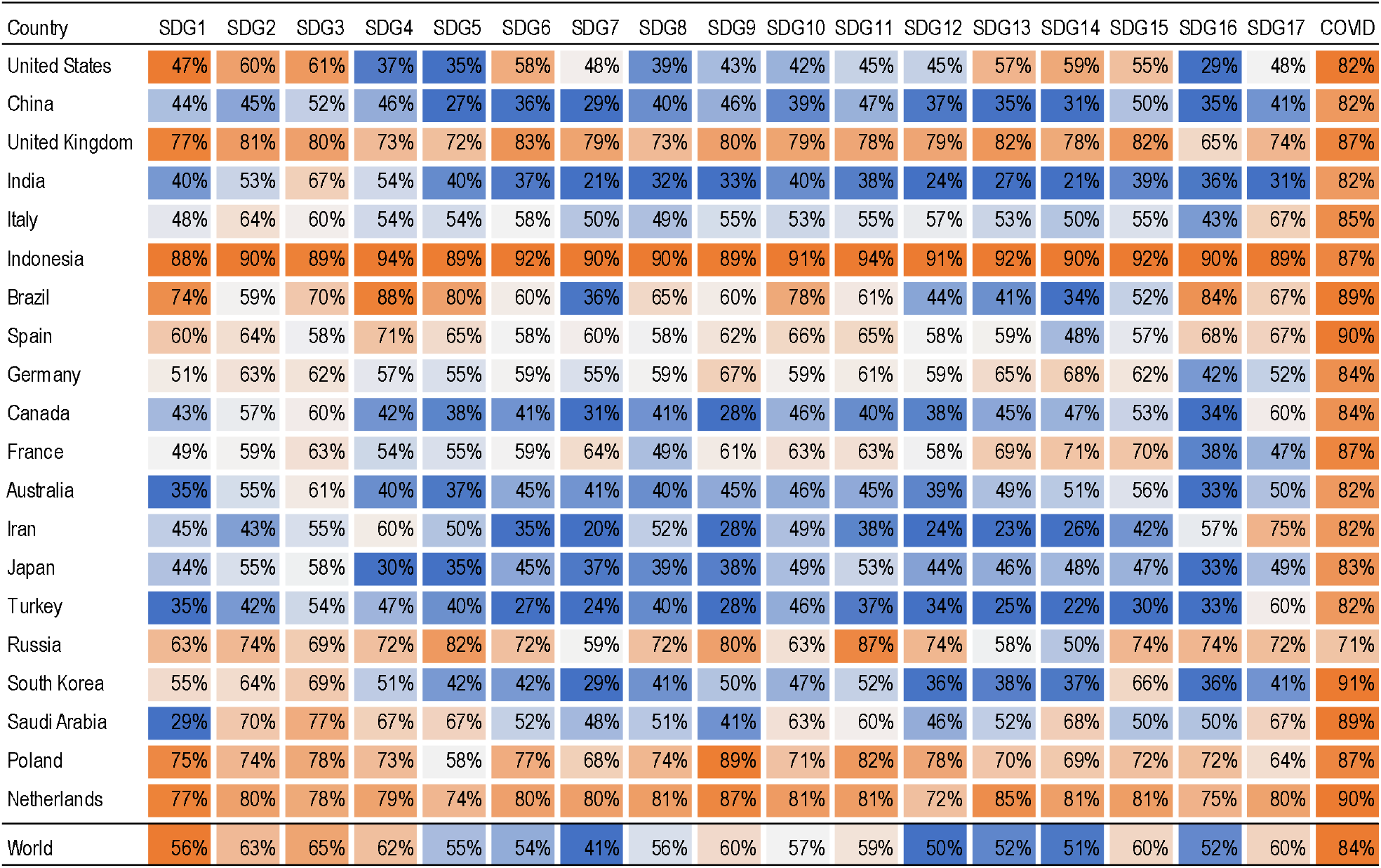
Percentage of Open Access papers, by SDG and country, 2017-2021. Dimensions.ai database.

## Footnotes

1 Wang, L. L., Lo, K., Chandrasekhar, Y., Reas, R., Yang, J., Eide, D., … & Kohlmeier, S. (2020). Cord-19: The COVID-19 open research dataset. ArXiv. Dataset available at: https://www.kaggle.com/datasets/allen-institute-for-ai/CORD-19-research-challenge

2 Armitage, C. S., Lorenz, M., & Mikki, S. (2020). Mapping scholarly publications related to the Sustainable Development Goals: Do independent bibliometric approaches get the same results?. Quantitative Science Studies, 1(3), 1092-1108.

3 Wastl, Jürgen, & Diwersy, Mario. (2019, December 19). Phase 1 and Phase 2 Summary of SDG Project by Springer Nature, VSNU/UKB, Digital Science. Zenodo. https://doi.org/10.5281/zenodo.3904447

4 Piwowar, H., Priem, J., Larivière, V., Alperin, J.P., Matthias, L., Norlander, B., Farley, A., West, J., Haustein, S. (2018) The State of OA: A large-scale analysis of the prevalence and impact of Open Access articles. PeerJ 6:e4375 https://doi.org/10.7717/peerj.4375

